# Biological age and brain age in midlife: relationship to multimorbidity and mental health

**DOI:** 10.1101/2022.09.26.509522

**Authors:** Fengqing Zhang, Hansoo Chang, Stacey M. Schaefer, Jiangtao Gou

**Affiliations:** Department of Psychological and Brain Sciences, Drexel University, Philadelphia, USA; Center for Healthy Minds, University of Wisconsin, Madison, Wisconsin, USA; Department of Mathematics and Statistics, Villanova University, Villanova, USA

**Keywords:** biological age, brain age, multimorbidity, mental health multimorbidity, machine learning

## Abstract

Multimorbidity, co-occurrence of two or more chronic conditions, is one of the top priorities in global health research and has emerged as the gold standard approach to study disease accumulation. As aging underlies the development of many chronic conditions, surrogate aging biomarkers are not disease-specific and capture health at the whole person level, having the potential to improve our understanding of multimorbidity. Biological age has been examined in recent years as a surrogate biomarker to capture the process of aging. However, relatively few studies have investigated the relationship between biological age and multimorbidity. More research is needed to quantify biological age using a broad range of biological markers and multimorbidity based on a comprehensive set of chronic conditions. Brain age estimated by neuroimaging data and machine learning models is another surrogate aging biomarker predictive of a wide range of health outcomes. Little is known about the relationship between brain age and multimorbidity. To answer these questions, our study investigates whether elevated biological age and accelerated brain age are associated with an increased risk of multimorbidity using a large dataset from the Midlife in the United States (MIDUS) Refresher study. Ensemble learning is utilized to combine multiple machine learning models to estimate biological age using a comprehensive set of biological markers. Brain age is obtained using convolutional neural networks and neuroimaging data. Our study is the first to examine the relationship between accelerated brain age and multimorbidity and presents the first effort to test whether sex moderates the relationship between these surrogate aging biomarkers and multimorbidity. Furthermore, it is the first attempt to explore how biological age and brain age are related to multimorbidity in mental health. Our findings hold the potential to advance the understanding of the accumulation of physical and mental health conditions, which may contribute to new strategies to improve the treatment of multimorbidity and detection of at-risk individuals.

## Introduction

Multimorbidity, coexistence of two or more chronic conditions in an individual, is increasingly common especially among older adults and poses substantial public health challenges (Suls, Green, & Davidson, 2016). Multimorbidity differs from a related term comorbidity in the sense that comorbidity has a primary condition of interest and examines the co-occurring of other conditions with this primary condition (Johnston, Crilly, Black, Prescott, & Mercer, 2019). Medical research has been primarily single disease focused, which considers one disease at a time and sometimes the comorbidity of a particular disease of interest (Suls et al., 2016). Multimorbidity offers an effective framework for examining the co-occurrence of multiple chronic conditions without the need to define a specific primary condition (Fabbri et al., 2015).

Aging is a universally experienced process, which underlies the development of most chronic conditions. The mechanism that drives the aging process may also drive multiple age-related chronic conditions. In this sense, aging represents a key risk factor for multimorbidity and a shared pathway across many chronic diseases (Fabbri et al., 2015; Liu et al., 2018). In recent years, biological age has been proposed as a surrogate biomarker to capture the process of aging (Klemera & Doubal, 2006; Levine, 2012). At the individual level, biological age can deviate from chronological age, with elevated biological age indicating higher risk for aging-related diseases (Wu et al., 2021). More importantly, this type of surrogate aging biomarker allows researchers to examine health at the whole person level, not unique to a specific health condition.

However, currently only three studies have investigated the relationship between biological age and multimorbidity. Biological age obtained from regression based models with a set of clinical markers was found to explain a significant amount of variability in multimorbidity defined based on five chronic diseases in middle-aged and older adults (Crimmins, Thyagarajan, Kim, Weir, & Faul, 2021). Another study reported a significant correlation between multimorbidity and a biological age metric estimated based on blood immune biomarkers in older adults (Sayed et al., 2021). Additionally, a strong association was reported between the number of diseases and a summary aging measure obtained by regressing the hazard of mortality on a set of clinical markers and chronological age in adult (Liu et al., 2018). More research is needed to examine biological age estimated from a broad range of clinical/biochemical markers and multimorbidity defined based on more categories of chronic conditions. In addition, a number of important methodological issues need to be clarified, such as how to address the estimation bias that biological age tends to be overestimated for younger subjects and underestimated for older subjects in regression-based models.

Brain age estimated by machine learning models and neuroimaging data is another surrogate biomarker on aging (X. Niu, Taylor, Shinohara, Kounios, & Zhang, 2022; Xin Niu, Zhang, Kounios, & Liang, 2020). Similar to biological age, brain age allows us to examine health as an integrative process, not specific to a particular health condition. Comparison of an individual’s brain age and chronological age can inform us whether an individual’s brain ages faster or slower than it should. Studies have shown that brain age is powerful in predicting a broad range of health outcomes, such as cognitive functioning, stroke, diabetes, Alzheimer’s disease, smoking, alcohol, and mortality (Cole, 2020; J. H. Cole et al., 2017; Wang et al., 2019). However, no study has examined the relationship between brain age and multimorbidity. In addition, it is unclear whether the relationship between these aging biomarkers (i.e., biological age, brain age) and multimorbidity is moderated by sex. Studies have reported that the prevalence of multimorbidity differs by gender, notably a higher rate in women (Marengoni et al., 2011). Understanding how the influence of biological age and brain age on multimorbidity differs by sex may allow us to develop more personalized strategies to prevent or delay multimorbidity.

To answer these questions, we first estimate biological age using machine learning models and a comprehensive set of clinical and biochemical markers including BMI, blood hemoglobin, and C-reactive protein. We then investigate whether elevated biological age is related to an increased risk of multimorbidity defined based on 13 different categories of chronic conditions. Then we examine whether accelerated brain age estimated by machine learning models and neuroimaging data is associated with an increased risk of multimorbidity. In addition, we test whether the relationship between multimorbidity and the surrogate aging biomarker (e.g., biological age and brain age) is moderated by sex. Lastly, multimorbidity in mental health has been relatively under-researched, despite its advantage in understanding clinical complexity in psychiatry (i.e., having multiple psychiatric and/or addictive disorders in an individual) (Bhalla, Stefanovics, & Rosenheck, 2018; Langan, Mercer, & Smith, 2013). To this end, we explore the relationship between these surrogate aging biomarkers and mental health multimorbidity. Findings from our study hold the potential to improve our understanding of multimorbidity and its major risk factor aging, which may shed the light on new strategies to improve the treatment and clinical management of multimorbidity.

## Methods

### MIDUS Refresher Study

We used data from the Midlife in the United States (MIDUS) Refresher study (Ryff et al., 2017). A national sample of 4085 adults, aged 25-74, were studied between 2011-2016. Participants recruited through RDD (random digit dialing) and a separate Black/African American sample from Milwaukee were included in the analyses. To estimate biological age, we used a subsample of subjects with biomarker assessments from the MIDUS Refresher Biomarker Project (n = 863). To obtain brain age, we used a subsample of individuals with neuroscience assessments from the MIDUS Refresher Neuroscience Project (n=138). Subjects in the Neuroscience Project were a subset of participants recruited in the Biomarker Project.

### Multimorbidity

We assessed multimorbidity using a dichotomous variable, indicating whether having two or more of the following conditions. Similar to other studies that examined multimorbidity using MIDUS data, we considered 13 different categories of chronic conditions including diabetes, asthma, hypertension, HIV or AIDS, tuberculosis, neurological disorders, stroke, ulcer, arthritis, ever had cancer, heart trouble, obesity, and/or high cholesterol levels (Friedman, Christ, & Mroczek, 2015; Shorey & Friedman, 2018). This dichotomous variable was coded as 0 if the subject had single or no condition and 1 if the subject had two or more of the 13 chronic conditions in the past 12 months. We computed multimorbidity for all subjects in the Biomarker Project. Additionally, we examined multimorbidity in mental health, a binary variable indicating whether having two or more of the psychiatric and addictive conditions including depression (Rottenberg, Devendorf, Panaite, Disabato, & Kashdan, 2019), anxiety disorder (Disabato et al., 2021), alcohol abuse (Ransome, Slopen, Karlsson, & Williams, 2017), and drug misuse (Kim et al., 2020).

### Biological measures

Based on studies that estimated biological age using clinical and biochemical markers (Belsky et al., 2015; Crimmins et al., 2021; Klemera & Doubal, 2006), we identified a set of fairly standard markers that are commonly collected in a clinical exam: BMI, waist-hip ratio, blood pressure (systolic), HDL cholesterol, LDL cholesterol, total cholesterol, triglycerides, HbA1C, blood fasting glucose, blood fasting IGF1 (insulin-like growth factor 1), C-reactive protein, creatinine, peak flow, blood serum MSD IL10, and bone specific alkaline phosphatase.

### Biological age

We built machine learning models to estimate biological age with the list of biological measures described above. To do so, we first identified a subset of healthy subjects without any of the physical and mental health conditions listed above (n=193, a mean age of 45.60, sd = 12.73, age range 25-74). The healthy cohort was randomly split in half with 50% of the data as a training set and the remaining 50% as an independent test set. We then built machine learning models with the list of biological measures to predict chronological age using the training set. In particular, we used ensemble learning to combine three machine learning models including support vector regression (Smola & Schölkopf, 2004), elastic net (Zou & Hastie, 2005), and Gaussian process regression (Rasmussen & Williams, 2006). Biological measures were standardized before running machine learning models. To tune the model parameters, 10-fold cross-validation was used within the training set. The trained model was then applied to the independent test set of healthy subjects to predict their chronological age. The predicted chronological age is the so-called biological age. The underlying assumption is that the biological age is on average the same as the chronological age for healthy subjects. In this sense, we would expect an accurate model to achieve a small mean absolute error (MAE) from a test set of healthy subjects. One important methodological issue that has not been made clear in the literature of biological age estimation is that a valid metric for evaluating model performance needs to be computed from an independent test set. This is because a model can easily overfit a training set. Another key methodological issue to note is that biological age can deviate from chronological age in disease groups. Thus, models for predicting biological age should be built with healthy subjects. Though model training should be done with a healthy cohort, the trained model for estimating biological age can be applied to disease groups. We also applied our trained model to predict biological age for subjects with at least one of the 13 categories of chronic conditions (n=630). We computed a difference score by subtracting chronological age from the estimated biological age, called biological age gap. A positive biological age gap means accelerated biological aging.

### Image acquisition and pre-processing

All structural scans were acquired using a 3T scanner (MR750 GE Healthcare, Waukesha, WI) with an 8-channel head coil. These data were derived from BRAVO T1-weighted structural images with TR = 8.2 ms, TE = 3.2 ms, flip angle = 12°, matrix = 256 × 256, FOV = 256 mm, slices = 160, slice thickness = 1 mm, and inversion time = 450 ms. Raw NIfTI files were minimally pre-processed using SPM12 including segmentation and normalization. FSL slicesdir (v. 5.0.11) was used to generate files for quality control in a web browser. The normalized images were loaded into R (v. 3.5.2) and vectorized. The mean image derived from the registration template for each tissue type was binarized with a threshold of 0.2 to create a mask. The resulting grey matter and white matter vectors were then masked and combined, which was used as the input of the brain age prediction model described below.

### Brain age

The estimates of brain age were computed using the BrainAgeR model published by Cole and colleagues (2017) (James H. Cole et al., 2017). In brief, convolutional neural networks were built using raw T1-weighted MRI data from a large set of healthy adults (n=2001, age range 18-90 years) to predict chronological age. This model was reported to predict chronological age accurately (Pearson’s correlation r = 0.94, MAE = 4.65 years). The predicted chronological age based on brain imaging data is the so-called brain-predicted age, or brain age in short. As discussed in previous publications from our group and others, brain age is often overestimated in younger individuals and underestimated in older individuals (James H. Cole et al., 2017; Liang, Zhang, & Niu, 2019; X. Niu et al., 2022; Xin Niu et al., 2020; Smith, Vidaurre, Alfaro-Almagro, Nichols, & Miller, 2019). This BrainAgeR model also corrects for age-related prediction bias by regressing brain age on chronological age. The trained model from Cole and colleagues (2017) (James H. Cole et al., 2017) was applied to raw T1-weighted MRI data collected in the Neuroscience Project (n=138) to obtain their brain age. Moreover, we calculated a difference score between brain age and chronological age, called brain age gap. A positive brain age gap indicates accelerated brain aging.

### Statistical analysis

To examine whether accelerated biological aging was associated with an increased risk of multimorbidity, we conducted logistic regression with biological age gap as the independent variable and multimorbidity as the outcome variable. In this analysis, we pooled the test set of healthy subject and the remaining subjects with at least one chronic condition together (total sample size n=726). We also controlled for chronological age, sex, race, and education. Race was coded as a binary variable, representing white and non-white. Education was categorized as three levels including high school or General Educational Development (GED), some college, and college degree or more. Similarly, logistic regression was used to examine whether advanced brain aging (i.e., a positive brain age gap) was related to an increased risk of multimorbidity. Since the brain age prediction model was trained using a different data set, we included all subjects from the MIDUS Refresher Neuroscience Project with structural MRI data available (n=127) in the logistic regression. The same set of covariates was controlled for in this analysis. In addition, we examined the potential moderating effect of sex on the relationship between multimorbidity and biological age gap. We also conducted separate moderation analyses to test whether the association between multimorbidity and brain age gap depended on sex. Lastly, we conducted exploratory analyses to examine the relationship between mental health multimorbidity and each of the two surrogate aging biomarkers (i.e., biological age gap, brain age gap).

## Results

### Sample characteristics

Descriptive statistics for the study sample is shown in Table 1. Subjects in this study aged between 25 and 76 with a mean age of 50.84 (sd = 13.41). The sample was primarily non-Hispanic white, and 52.1% were female. Of the study sample, 52.2% had a college degree or more and 42.8% had two or more chronic conditions (i.e., multimorbidity). The proportion of mental health multimorbidity was 9.7%.

**Table 1.** Subject characteristics, the MIDUS Refresher Biomarker Project (n=863)

| Variable | Mean (SD) | Range | Percent |
| --- | --- | --- | --- |
| Age | 50.84 (13.41) | 25-76 |  |
| Sex (% female) |  |  | 52.1% |
| Race (% non-White) |  |  | 29.4% |
| Education |  |  |  |
| High school or GED |  |  | 17.3% |
| Some college |  |  | 30.5% |
| College or more |  |  | 52.2% |
| Number of chronic conditions | 1.58 (1.56) | 0-8 |  |
| Multimorbidity (% yes) |  |  | 42.8% |
| Number of mental health conditions | 0.42 (0.74) | 0-4 |  |
| Mental health multimorbidity (% yes) |  |  | 9.7% |

### Biological age

In the training set, biological age estimated by the machine learning model was correlated with chronological age (r = 0.81, MAE = 7.09). As shown in Figure 1A, biological age was overestimated for younger subjects and underestimated for older subjects. Due to this systematic difference between the estimated biological age and chronological age, biological age gap was negatively associated with chronological age (Figure 1B). However, we expect the biological age gap is on average zero among healthy subjects. To account for this systematic negative association between biological age gap and chronological age, we regressed biological age gap on chronological age and computed the residualized/adjusted biological age gap (i.e., bias correction step). As shown in Figure 1C, the adjusted biological age gap was not associated with chronological age and was on average very close to zero. When investigating biological age or biological age gap as a potential biomarker for aging, it is essential to account for this systematic bias by either computing the residualized biological age (gap) or controlling for chronological age. Otherwise, the effect of biological age (gap) can be confounded by chronological age.

**Figure 1.**
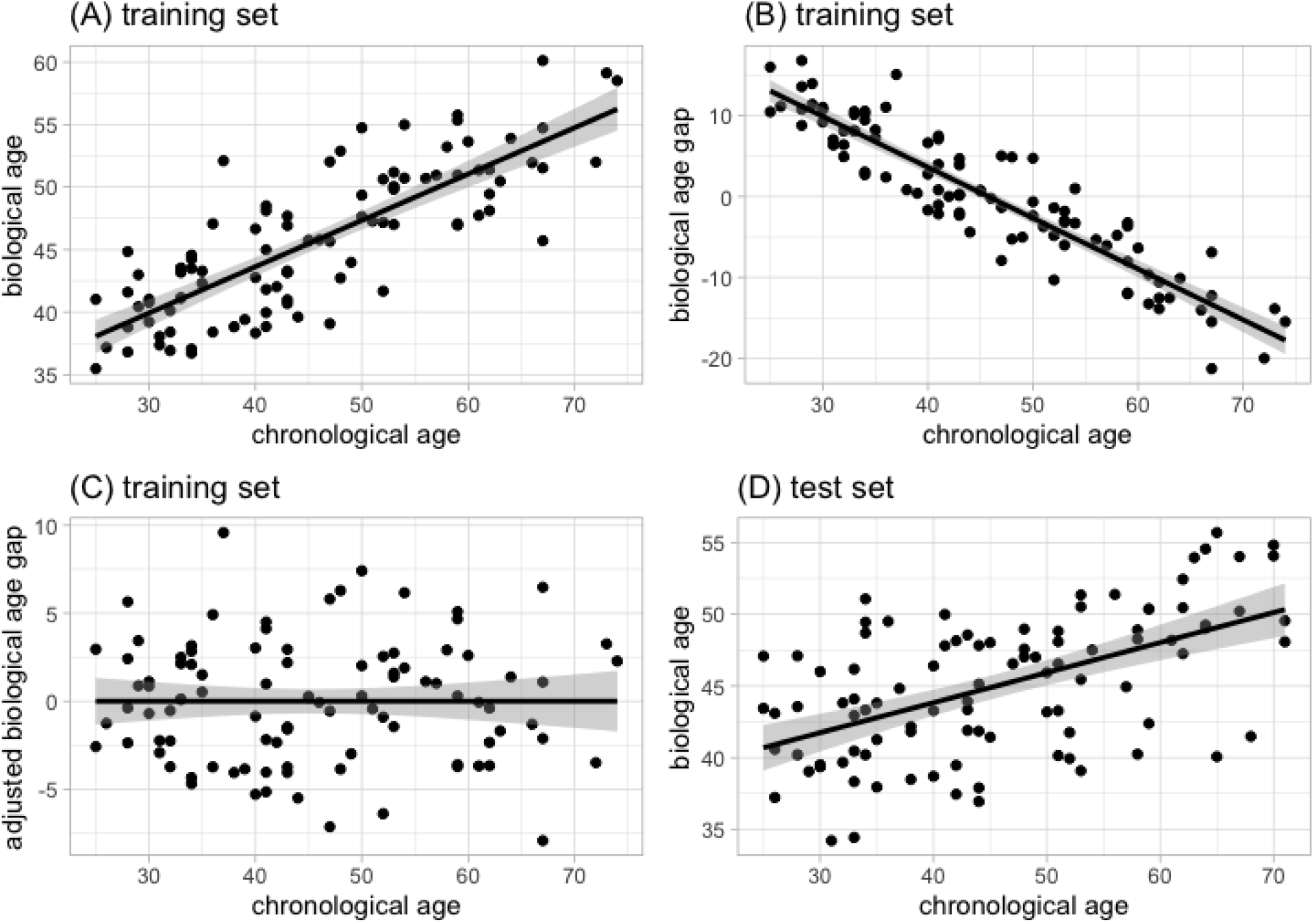
Scatterplots of biological age/(adjusted) biological age gap (y-axis) and chronological age (x-axis) with fitted regression lines and 95% confidence bands for the training and test sets of healthy adults.

The trained machine learning model was then applied to a test set of healthy adults to evaluate model performance. The systematic difference in the estimated biological age and chronological age was also observed in the test set (Figure 1D). The model performance on the test set (r = 0.54, MAE = 9.08) was worse than what was achieved on the training set. This highlights the importance of reporting model performance from both the training and test sets. It is also worth noting that a high correlation coefficient and a small MAE imply better prediction performance only when we examine a cohort of healthy subjects.

Feature importance value was computed from the machine learning model to rank the biological markers in terms of their contribution for estimating biological age (Figure 2). The top predictors were total cholesterol, blood fasting IGF1 (insulin-like growth factor 1), bone specific alkaline phosphatase, and blood pressure. Creatinine was ranked as the least informative biological marker for estimating biological age.

**Figure 2.**
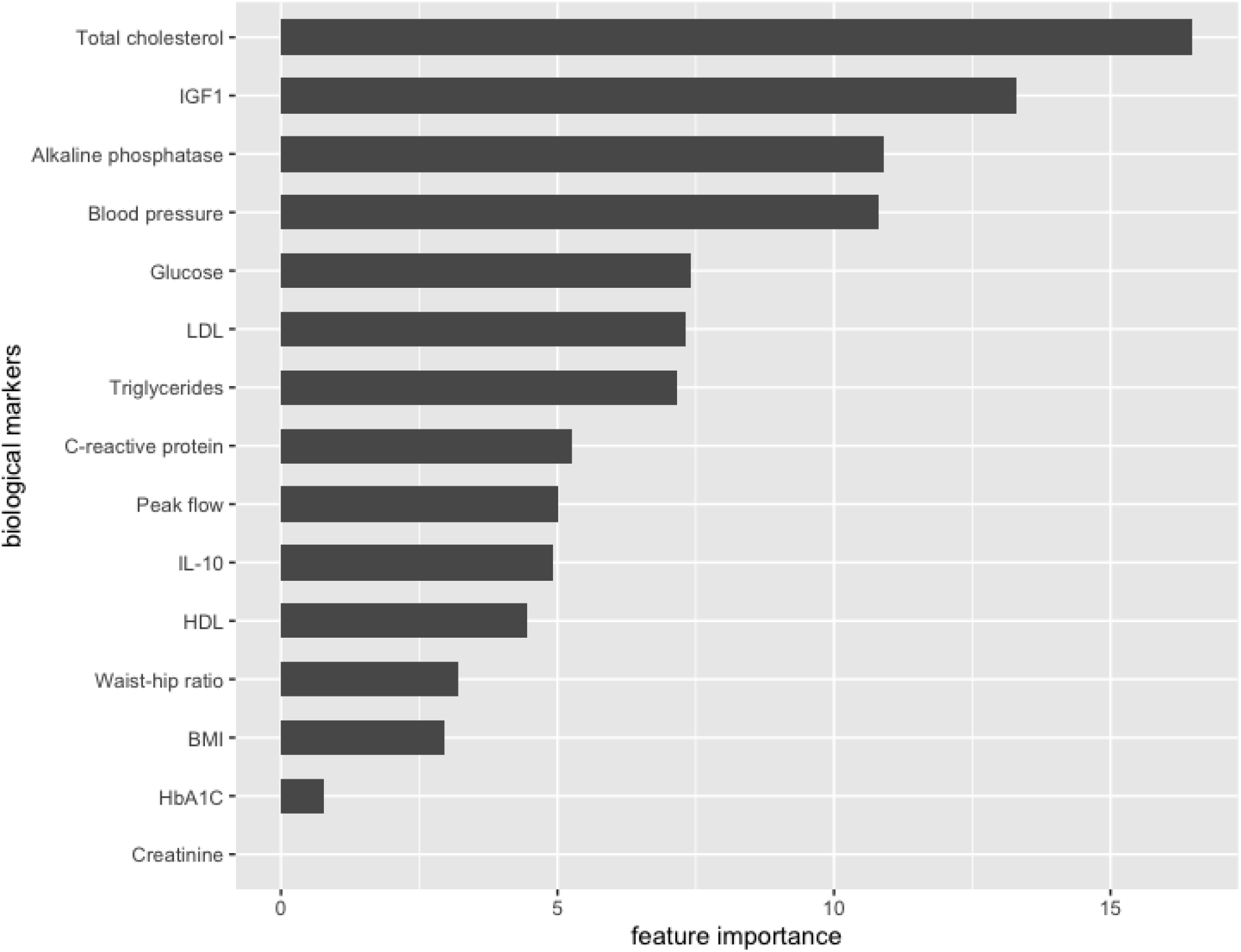
Feature importance yielded by the machine learning models for the list of biological markers used to estimate biological age.

### Brain age

The BrainAgeR model trained with a large set of healthy adults in Cole and colleagues (2017) was applied to the MIDUS Refresher Study Neuroscience Project and achieved good model performance (r = 0.79, MAE = 6.1). Similar to what we observed for biological age, brain age was overestimated in younger subjects and underestimated in older subjects (Figure 3 left panel). This systematic bias has been discussed in our previous publications(Liang et al., 2019; Xin Niu et al., 2020). Though a regression-based bias correction step was conducted automatically by the BrainAgeR model to account for a statistical dependency on chronological age(James H. Cole et al., 2017), we still observed some overestimation and underestimation in brain age (Figure 3 right panel). This again illustrates the importance of controlling for chronological age when using brain age (gap) as a potential biomarker for aging or other health conditions.

**Figure 3.**
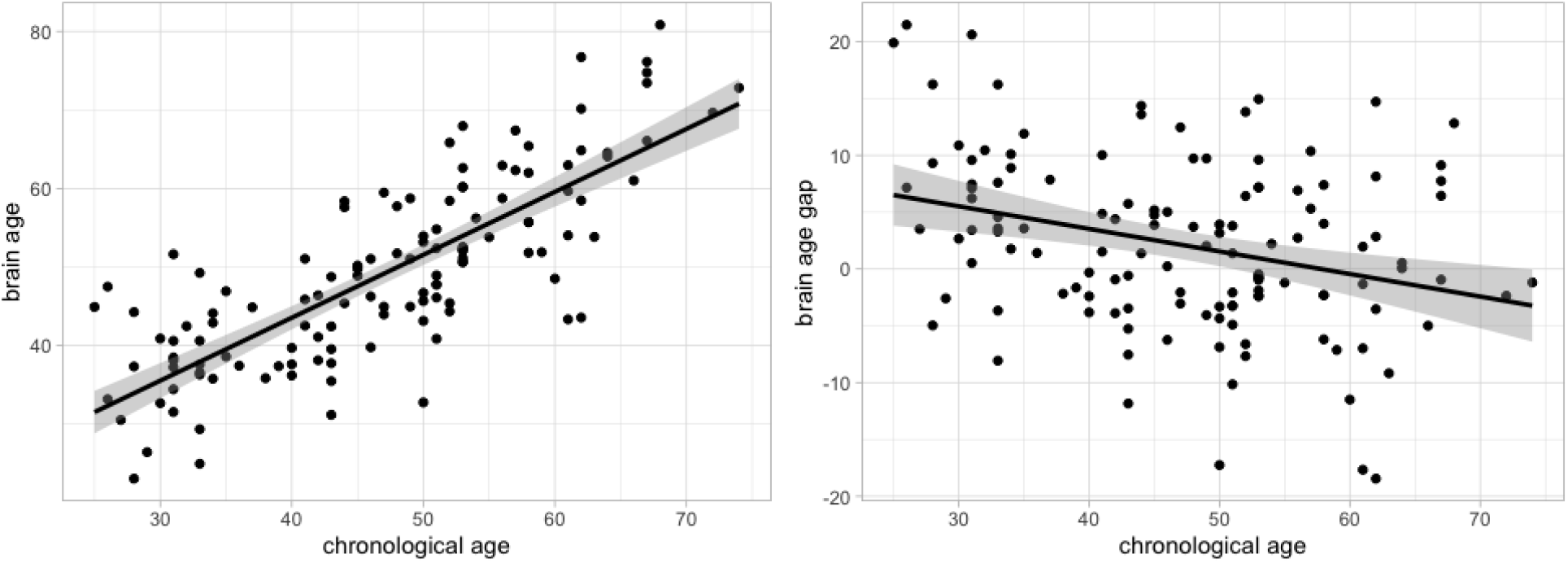
Scatterplots of brain age/brain age gap (y-axis) and chronological age (x-axis) with fitted regression lines and 95% confidence bands for the test set.

### Association with multimorbidity

As shown in Table 2, results from logistic regression found that a larger biological age gap was significantly associated with an increased risk of multimorbidity after controlling for sex, chronological age, race, and education (b = 0.06, p = 0.007, OR = 1.07, 95% CI [1.02, 1.12]). Among the covariates, higher chronological age (b = 0.13, p < 0.001, OR = 1.14, 95% CI [1.09, 1.19]) and being non-white (b = 0.45, p = 0.022, OR = 1.57, 95% CI [1.07, 2.30]) were significantly related to an increased risk of multimorbidity. Having a college degree or more was negatively associated with the risk of multimorbidity (b = −0.76, p < 0.001, OR = 0.48, 95% CI [0.30, 0.73]).

**Table 2.** Results from the logistic regression models for examining the association between biological age gap and multimorbidity and for examining the moderating effect of sex. Multimorbidity is defined based on 13 different categories of chronic conditions.

|  | Multimorbidity |  | Multimorbidity<br>(with sex as a moderator) |  |
| --- | --- | --- | --- | --- |
|  | b | P value | b | P value |
| Biological age gap | 0.06 | **0.007** | 0.06 | **0.011** |
| Sex (female) | 0.03 | 0.870 | 0.04 | 0.822 |
| Age | 0.13 | < **0.001** | 0.13 | < **0.001** |
| Race (non-white) | 0.45 | **0.022** | 0.45 | **0.022** |
| Education (some college) | -0.17 | 0.475 | -0.17 | 0.478 |
| Education (college or more) | -0.76 | < **0.001** | -0.76 | **0.001** |
| Biological age gap * Sex | --- | --- | 0.003 | 0.850 |

Findings on the association between brain age gap and multimorbidity were summarized in Table 3. Brain age gap was not significantly associated with the risk of multimorbidity (b = −0.02, p = 0.551, OR = 0.98, 95% CI [0.92, 1.04]). Among the covariates, chronological age was positively associated with the risk of multimorbidity (b = 0.11, p < 0.001, OR = 1.11, 95% CI [1.07, 1.17]) while having a college degree or more was negatively related to the risk of multimorbidity (b = −1.36, p = 0.021, OR = 0.26, 95% CI [0.08, 0.79]).

**Table 3.** Results from the logistic regression models for examining the association between brain age gap and multimorbidity and for examining the moderating effect of sex. Multimorbidity is quantified using 13 different categories of chronic conditions.

|  | Multimorbidity |  | Multimorbidity<br>(with sex as a moderator) |  |
| --- | --- | --- | --- | --- |
|  | b | P value | b | P value |
| Brain age gap | -0.02 | 0.551 | 0.07 | 0.129 |
| Sex (female) | 0.27 | 0.554 | 0.59 | 0.236 |
| Age | 0.11 | < **0.001** | 0.11 | < **0.001** |
| Race (non-white) | -0.03 | 0.951 | -0.26 | 0.608 |
| Education (some college) | -0.72 | 0.216 | -1.04 | 0.088 |
| Education (college or more) | -1.36 | **0.021** | -1.76 | **0.006** |
| Brain age gap * Sex | --- | --- | -0.19 | **0.009** |

### Moderating effect of sex

The effect of biological age gap on the risk of multimorbidity was not significantly moderated by sex (b = 0.003, p = 0.850, OR = 1.00, 95% CI [0.97, 1.03]). Including the interaction term between biological age gap and sex in the logistic regression did not change the pattern and significance of other variables (see the last two columns of Table 2). In contrast, the brain age gap by sex interaction term was statistically significant and was negatively associated with the risk of multimorbidity (b = −0.19, p = 0. 009, OR = 0.83, 95% CI [0.71, 0.95]). This suggests that the effect of brain age gap on the risk of multimorbidity depended on sex and was weaker among females compared to males. The inclusion of the interaction term did not change the pattern and significance of other variables in the logistic regression model (refer to the last two columns of Table 3).

### Association with mental health multimorbidity

In the exploratory analyses, we found that biological age gap was positively associated with the risk of mental health multimorbidity after controlling for sex, chronological age, race, and education (b = 0.07, p = 0.046, OR = 1.07, 95% CI [1.00, 1.15]). Among the covariates, having some college education was negatively related to the risk of mental health multimorbidity (b = −0.66, p = 0.031, OR = 0.52, 95% CI [0.28, 0.94]). Additionally, having a college degree or more was associated with a decreased risk of mental health multimorbidity (b = −1.17, p < 0.001, OR = 0.31, 95% CI [0.17, 0.57]).

Similarly, brain age gap was found to be positively related to the risk of mental health multimorbidity after controlling for sex, chronological age, race, and education (b = 0.10, p = 0.038, OR = 1.10, 95% CI [1.01, 1.22]). All other covariates were not statistically significant. This is likely due to the fact that the sample size of the Neuroscience Project is much smaller compared to the Biomarker Project (see Methods Section).

## Discussion

In this study, we estimated biological age using machine learning models and a comprehensive set of biological markers collected from a large number of adults in the MIDUS Refresher sample. We showed that elevated biological age was associated with an increased risk of multimorbidity defined based on 13 different categories of chronic conditions. In addition, it is the first study to examine the relationship between accelerated brain age and multimorbidity. We also investigated whether the relationship between the surrogate aging biomarkers and multimorbidity was moderated by sex. Though brain age gap was not associated with the risk of multimorbidity, the interaction between brain age gap and sex was significantly negatively related to the risk of multimorbidity, suggesting the effect of brain age gap on risk of multimorbidity was weaker among females compared to males. Lastly, in our exploratory analyses, we found elevated biological age and accelerated brain age were both associated with an increased risk of mental health multimorbidity. These findings are helpful to improve our understanding of the accumulation of physical and mental health conditions in an individual and sex-related differences in aging.

Though varied in the methods of estimating biological age and quantifying multimorbidity, our study obtained findings consistent with previous studies that elevated biological age was related to an increased risk of multimorbidity (Crimmins et al., 2021; Liu et al., 2018; Sayed et al., 2021). This points to the value of investigating biological aging as an underlying mechanism for multimorbidity and a shared pathway for different chronic conditions. Early detection of accelerated biological aging before the onset of chronic conditions holds the potential for disease prevention. Cole 2020 reported that advanced brain aging was associated with hypertension, diabetes, and stroke when testing with 14,701 individuals from UK Biobank (Cole, 2020). However, in our study brain age was not found to be significantly associated with multimorbidity. The relatively small sample size in the Neuroscience Project could limit the statistical power to detect the effect. As our study is the first attempt to explore the association between brain age and multimorbidity, future studies with larger sample sizes are needed to validate the findings.

We identified a set of biological measures, which were found to influence biological age that was positively associated with the risk of multimorbidity. Our top ranked biological markers (i.e., feature importance >= 5) for estimating biological age including total cholesterol, IGF1, alkaline phosphatase, glucose, C-reactive protein, and peak flow have also been reported to be predictive of multimorbidity (Crimmins et al., 2021). Systolic blood pressure has been identified as a risk factor for death and disease (Liu et al., 2018; Port, Demer, Jennrich, Walter, & Garfinkel, 2000). In addition to total cholesterol, LDL and triglycerides were also found to be important for the quantification of biological aging (Belsky et al., 2015). These identified biological markers are modifiable factors, which may shed the light on new approaches to improve the treatment of multimorbidity. This also points to the need of collecting these biological measures during routine medical visits, which may assist with the detection of at-risk individuals.

The negative association between brain age gap and chronological age has been thoroughly investigated in prior work published by our group (Liang et al., 2019; X. Niu et al., 2022; Xin Niu et al., 2020) as well as other studies (James H. Cole et al., 2017; Smith et al., 2019). The reason for this systematic bias can be explained by regression to the mean (Liang et al., 2019). However, this issue has not been formally reported in the literature of biological age estimation. Thus, we presented the pattern between biological age gap and chronological age using the MIDUS Refresher sample and highlighted the importance of controlling for the confounding effect of chronological age. Additionally, our study also demonstrated the need of evaluating model performance using an independent test set as statistical models can easily overfit a training dataset. Because biological age can deviate from chronological in disease groups, correlation coefficient and MAE are only meaningful metrics for evaluating model performance when examining healthy subjects. This speaks to the value of training/calibrating biological age estimation model only using health subjects. Presenting these methodological issues in our study is helpful to guide the design and analysis of future studies on biological age and brain age.

Our study found that the two-way interaction between brain age and sex was negatively associated with the risk of multimorbidity. This suggests the potential gender difference in how brain aging is related to the accumulation of multiple chronic conditions. Previous study on brain aging reported on average younger brain in women throughout adulthood compared to men of the same age (Goyal et al., 2019). Predictors of brain age were also found to be sex-specific, highlighting the value of sex-specific analyses (Sanford et al., 2022). Further research is needed on examining sex-specific risk and protective factors that influence brain aging and disease accumulation.

Multimorbidity in mental health has been relatively under-investigated, despite its strength in capturing complex clinical representations of psychiatric disorders (Bhalla et al., 2018; Langan et al., 2013). Our study presents the first attempt to investigate how biological age and brain age are related to mental health multimorbidity. As hypothesized, we found elevated biological age and accelerated brain age were associated with an increased risk of mental health multimorbidity. Our findings are consistent with previous studies that found accelerated brain aging in alcohol use (Amen, Egan, Meysami, Raji, & George, 2018; Cole, 2020), cannabis use (Amen et al., 2018), anxiety (Amen et al., 2018), and depression (X. Niu et al., 2022). In addition, accelerated biological aging has been reported in substance use (Bachi, Sierra, Volkow, Goldstein, & Alia-Klein, 2017) and alcohol abuse (Piniewska-Róg et al., 2021). Some biological age indicators (e.g., lung function, telomere length) have also been found to be altered in depression and anxiety disorder (Han et al., 2019). It will be interesting to examine the relationship between these surrogate aging biomarkers and mental health multimorbidity using data with larger samples and more categories of mental health conditions.

Because of our interest in examining brain age estimated by neuroimaging data, we chose the MIDUS refresher sample in this study and thus our findings are limited to cross-sectional association. Future research can benefit from longitudinal data to investigate whether biological age and brain age predict multimorbidity and mental health multimorbidity prospectively. The longitudinal nature of the MIDUS study will allow such future investigations. Furthermore, the estimated brain age was obtained based on structural MRI data. It will be interesting to test whether brain age estimated by multimodal neuroimaging data shows a stronger association with multimorbidity and mental health multimorbidity. In addition, our study defined the primary outcome multimorbidity based on 13 different categories of chronic conditions and explored mental health multimorbidity based on measures available in the MIDUS refresher sample. Alternative ways of defining multimorbidity (e.g., whether the duration and severity of chronic conditions are considered) and physical-mental multimorbidity patterns need to be examined in future research. Integrating other lifestyle factors such as diet and physical activity in the relationship between aging and multimorbidity may merit future research.

## References

Amen, D. G., Egan, S., Meysami, S., Raji, C. A., & George, N. (2018). Patterns of regional cerebral blood flow as a function of age throughout the lifespan. Journal of Alzheimer’s disease, 65(4), 1087–1092.

Bachi, K., Sierra, S., Volkow, N. D., Goldstein, R. Z., & Alia-Klein, N. (2017). Is biological aging accelerated in drug addiction? Current Opinion in Behavioral Sciences, 13, 34–39. doi:10.1016/j.cobeha.2016.09.007

Belsky, D. W., Caspi, A., Houts, R., Cohen, H. J., Corcoran, D. L., Danese, A., … Moffitt, T. E. (2015). Quantification of biological aging in young adults. Proceedings of the National Academy of Sciences, 112(30). doi:10.1073/pnas.1506264112

Bhalla, I. P., Stefanovics, E. A., & Rosenheck, R. A. (2018). Mental health multimorbidity and poor quality of life in patients with schizophrenia. Schizophrenia Research, 201, 39–45. doi:10.1016/j.schres.2018.04.035

Cole, J. H. (2020). Multimodality neuroimaging brain-age in UK biobank: relationship to biomedical, lifestyle, and cognitive factors. Neurobiology of Aging, 92, 34–42. doi:10.1016/j.neurobiolaging.2020.03.014

Cole, J. H., Poudel, R. P. K., Tsagkrasoulis, D., Caan, M. W. A., Steves, C., Spector, T. D., & Montana, G. (2017). Predicting brain age with deep learning from raw imaging data results in a reliable and heritable biomarker. NeuroImage, 163, 115–124. doi:10.1016/j.neuroimage.2017.07.059

Cole, J. H., Ritchie, S. J., Bastin, M. E., Valdés Hernández, M. C., Muñoz Maniega, S., Royle, N., … Deary, I. J. (2017). Brain age predicts mortality. Molecular Psychiatry, 23(5), 1385–1392. doi:10.1038/mp.2017.62

Crimmins, E. M., Thyagarajan, B., Kim, J. K., Weir, D., & Faul, J. (2021). Quest for a summary measure of biological age: the health and retirement study. GeroScience, 43(1), 395–408. doi:10.1007/s11357-021-00325-1

Disabato, D. J., Kashdan, T. B., Doorley, J. D., Kelso, K. C., Volgenau, K. M., Devendorf, A. R., & Rottenberg, J. (2021). Optimal well-being in the aftermath of anxiety disorders: A 10-year longitudinal investigation. Journal of affective disorders, 291, 110–117. doi:10.1016/j.jad.2021.05.009

Fabbri, E., Zoli, M., Gonzalez-Freire, M., Salive, M. E., Studenski, S. A., & Ferrucci, L. (2015). Aging and multimorbidity: new tasks, priorities, and frontiers for integrated gerontological and clinical research. Journal of the American Medical Directors Association, 16(8), 640–647. doi:10.1016/j.jamda.2015.03.013

Friedman, E. M., Christ, S. L., & Mroczek, D. K. (2015). Inflammation partially mediates the association of multimorbidity and functional limitations in a national sample of middleaged and older adults. Journal of Aging and Health, 27(5), 843–863. doi:10.1177/0898264315569453

Goyal, M. S., Blazey, T. M., Su, Y., Couture, L. E., Durbin, T. J., Bateman, R. J., … Vlassenko, A. G. (2019). Persistent metabolic youth in the aging female brain. Proceedings of the National Academy of Sciences, 116(8), 3251–3255. doi:10.1073/pnas.1815917116

Han, L. K. M., Verhoeven, J. E., Tyrka, A. R., Penninx, B. W. J. H., Wolkowitz, O. M., Månsson, K. N. T., … Picard, M. (2019). Accelerating research on biological aging and mental health: Current challenges and future directions. Psychoneuroendocrinology, 106, 293–311. doi:10.1016/j.psyneuen.2019.04.004

Johnston, M. C., Crilly, M., Black, C., Prescott, G. J., & Mercer, S. W. (2019). Defining and measuring multimorbidity: a systematic review of systematic reviews. European Journal of Public Health, 29(1), 182–189. doi:10.1093/eurpub/cky098

Kim, E. S., Ryff, C., Hassett, A., Brummett, C., Yeh, C., & Strecher, V. (2020). Sense of purpose in life and likelihood of future illicit drug use or prescription dedication misuse. Psychosomatic Medicine, 82(7), 715–721. doi:10.1097/psy.0000000000000842

Klemera, P., & Doubal, S. (2006). A new approach to the concept and computation of biological age. Mechanisms of Ageing and Development, 127(3), 240–248.

Langan, J., Mercer, S. W., & Smith, D. J. (2013). Multimorbidity and mental health: can psychiatry rise to the challenge? British Journal of Psychiatry, 202(6), 391–393. doi:10.1192/bjp.bp.112.123943

Levine, M. E. (2012). Modeling the rate of senescence: can estimated biological age predict mortality more accurately than chronological age? The Journals of Gerontology Series A: Biological Sciences and Medical Sciences, 68(6), 667–674. doi:10.1093/gerona/gls233

Liang, H., Zhang, F., & Niu, X. (2019). Investigating systematic bias in brain age estimation with application to post traumatic stress disorders. Human Brain Mapping, 40(11), 3143–3152. doi:10.1002/hbm.24588

Liu, Z., Kuo, P.-L., Horvath, S., Crimmins, E., Ferrucci, L., & Levine, M. (2018). A new aging measure captures morbidity and mortality risk across diverse subpopulations from NHANES IV: A cohort study. PLOS Medicine, 15(12), e1002718. doi:10.1371/journal.pmed.1002718

Marengoni, A., Angleman, S., Melis, R., Mangialasche, F., Karp, A., Garmen, A., … Fratiglioni, L. (2011). Aging with multimorbidity: A systematic review of the literature. Ageing Research Reviews, 10(4), 430–439. doi:10.1016/j.arr.2011.03.003

Niu, X., Taylor, A., Shinohara, R. T., Kounios, J., & Zhang, F. (2022). Multidimensional brainage prediction reveals altered brain developmental trajectory in psychiatric disorders. Cerebral Cortex, bhab530. Advance online publication.

Niu, X., Zhang, F., Kounios, J., & Liang, H. (2020). Improved prediction of brain age using multimodal neuroimaging data. Human Brain Mapping, 41(6), 1626–1643. doi:10.1002/hbm.24899

Piniewska-Róg, D., Heidegger, A., Pośpiech, E., Xavier, C., Pisarek, A., Jarosz, A., … Branicki, W. (2021). Impact of excessive alcohol abuse on age prediction using the VISAGE enhanced tool for epigenetic age estimation in blood. International Journal of Legal Medicine, 135(6), 2209–2219. doi:10.1007/s00414-021-02665-1

Port, S., Demer, L., Jennrich, R., Walter, D., & Garfinkel, A. (2000). Systolic blood pressure and mortality. The Lancet, 355(9199), 175–180. doi:10.1016/s0140-6736(99)07051-8

Ransome, Y., Slopen, N., Karlsson, O., & Williams, D. R. (2017). Elevated inflammation in association with alcohol abuse among Blacks but not Whites: results from the MIDUS biomarker study. Journal of Behavioral Medicine, 41(3), 374–384. doi:10.1007/s10865-017-9905-4

Rasmussen, C. E., & Williams, C. K. I. (2006). Gaussian processes for machine learning: The MIT Press.

Rottenberg, J., Devendorf, A. R., Panaite, V., Disabato, D. J., & Kashdan, T. B. (2019). Optimal Well-Being after Major Depression. Clinical Psychological Science, 7(3), 621–627. doi:10.1177/2167702618812708

Ryff, C., Almeida, D. M., Ayanian, J. Z., Binkley, N., Carr, D. S., Coe, C., … Williams, D. R. (2017). Midlife in the United States (MIDUS Refresher 1), 2011–2014.

Sanford, N., Ge, R., Antoniades, M., Modabbernia, A., Haas, S. S., Whalley, H. C., … Frangou, S. (2022). Sex differences in predictors and regional patterns of brain age gap estimates. Human Brain Mapping. doi:10.1002/hbm.25983

Sayed, N., Huang, Y., Nguyen, K., Krejciova-Rajaniemi, Z., Grawe, A. P., Gao, T., … Furman, D. (2021). An inflammatory aging clock (iAge) based on deep learning tracks multimorbidity, immunosenescence, frailty and cardiovascular aging. Nature Aging, 1(7), 598–615. doi:10.1038/s43587-021-00082-y

Shorey, C., & Friedman, E. (2018). Multimorbidity and cognitive decline in a national sample of aging adults. Innovation in Aging, 2, 505–506.

Smith, S. M., Vidaurre, D., Alfaro-Almagro, F., Nichols, T. E., & Miller, K. L. (2019). Estimation of brain age delta from brain imaging. NeuroImage, 200, 528–539. doi:10.1016/j.neuroimage.2019.06.017

Smola, A. J., & Schölkopf, B. (2004). A tutorial on support vector regression. Statistics and Computing, 14(3), 199–222.

Suls, J., Green, P. A., & Davidson, K. W. (2016). A biobehavioral framework to address the emerging challenge of multimorbidity. Psychosomatic Medicine, 78(3), 281–289. doi:10.1097/psy.0000000000000294

Wang, J., Knol, M. J., Tiulpin, A., Dubost, F., de Bruijne, M., Vernooij, M. W., … Roshchupkin, G. V. (2019). Gray matter age prediction as a biomarker for risk of dementia. Proceedings of the National Academy of Sciences, 116(42) 21213–21218. doi:10.1073/pnas.1902376116

Wu, J. W., Yaqub, A., Ma, Y., Koudstaal, W., Hofman, A., Ikram, M. A., … Goudsmit, J. (2021). Biological age in healthy elderly predicts aging-related diseases including dementia. Scientific Reports, 11(1). doi:10.1038/s41598-021-95425-5

Zou, H., & Hastie, T. (2005). Regularization and variable selection via the elastic net. Journal of the Royal Statistical Society. Series B (Statistical Methodology), 67(2), 301–320.

